# The School Attachment Monitor - a novel computational tool for assessment of attachment in middle childhood

**DOI:** 10.1101/2020.09.24.311258

**Authors:** Maki Rooksby, Simona Di Folco, Mohammad Tayarani, Dong-Bach Vo, Rui Huan, Alessandro Vinciarelli, Stephen A. Brewster, Helen Minnis

## Abstract

**Background:** Attachment research has been limited by the lack of quick and easy measures. We report development and validation of the School Attachment Monitor (SAM), a novel measure for largescale assessment of attachment in children aged 5-9, in the general population. SAM offers automatic presentation, on computer, of story-stems based on the Manchester Child Attachment Story Task (MCAST), without the need for trained administrators. SAM is delivered by novel software which interacts with child participants, starting with warm-up activities to familiarise them with the task. Children’s story completion is video recorded and augmented by ‘smart dolls’ that the child can hold and manipulate, with movement sensors for data collection. The design of SAM was informed by children of users’ age range to establish their task understanding and incorporate their innovative ideas for improving SAM software.

**Methods:** 130 5-9 year old children were recruited from mainstream primary schools. In Phase 1, sixty-one children completed both SAM and MCAST. Inter-rater reliability and rating concordance was compared between SAM and MCAST. In Phase 2, a further 44 children completed SAM complete and, including those children completing SAM in Phase 1 (total n=105), a machine learning algorithm was developed using a “majority vote” procedure where, for each child, 500 non-overlapping video frames contribute to the decision.

**Results:** Using manual rating, SAM-MCAST concordance was excellent (89% secure versus insecure; 97% organised versus disorganised; 86% four-way). Comparison of human ratings of SAM versus the machine learning algorithm showed over 80% concordance.

**Conclusions:** We have developed a new tool for measuring attachment at the population level, which has good reliability compared to a gold-standard attachment measure and has the potential for automatic rating – opening the door to measurement of attachment in large populations.

## Introduction

Attachment is a fundamental element of human psychological development, examined in thousands of studies worldwide. Yet this wealth of attachment research has failed to influence clinical practice on a day to day basis because rigorous attachment measures, developed for research purposes, are not appropriate to conduct and rate meaningfully in busy clinics (1). We aim, in this study, to develop and test an attachment measure that is quicker and easier to administer and rate than the existing gold standard tools.

These gold standard attachment measures provide a standardised, replicable protocol that presents the child with a series of stressful scenarios in the presence of the actual parent/caregiver or a representation of the child and parent/caregiver (e.g. using drawings or doll play). Determination of the attachment classification is achieved by direct observation of actual or represented child and parent/caregiver behaviours, and rigorous training is required to ensure these observations are valid and reliable (1). The Strange Situation Procedure (SSP) (2) which is one of the gold standard measures, used in many hundreds of studies, is most valid when used at a child age of 12 to 18 months and can also be used in the pre-school years (1). The Manchester Child Attachment story task – most valid when used in children aged 4 to 9 years of age – has been used in more than 25 studies across 9 countries and has had rigorous psychometric testing with adequate inter-rater (3), test-retest reliability and criterion validity (1).

However, measuring attachment is challenging and burdensome. For example, administering the SSP in clinical practice takes approximately an hour of two staff members’ time: a specifically trained administrator and a “stranger” plus at least an additional hour rating time. Training to administer and rate the SSP requires a two-week (usually US-based) course, then rigorous reliability training to reach at least 80% agreement with expert raters. Thereafter, high quality research labs achieve four-way attachment classifications of variable reliability (Cohen’s Kappas 0.49 to 0.93) (4, 5). So despite gold standard attachment measures being recommended (e.g. by the UK National Institute for Health and Care Excellence (6)), their use is rarely feasible in clinical practice.

Despite the large research database on attachment that has accrued since the 1970s, important questions remain unanswered e.g. how genetics and environment interact in the development of attachment relationships across the lifespan (7) and the potential that secure attachment could act as a “susceptibility factor”, encouraging children to explore the environment, resulting in either positive or negative outcomes depending on the risks and protective factors they encounter (8). To fully examine these important but complex questions, much larger samples will be necessary - in the order of the tens of thousands of participants (9).

Over a decade ago, we set out to try and address the challenge of finding a quicker and easier, yet valid, way to assess attachment patterns in early to middle childhood. First, we found that no quick and easy childhood measures of attachment existed (10), similarly to more recent systematic reviews (1, 11). Rapid and easy measures of attachment patterns in young children do not exist for good reasons: attachment is a relational process involving measurement of the child’s attachment behaviours, the caregiver’s caregiving behaviours and the degree to which the child’s distress is assuaged (3). Because attachment behaviours are only activated when the child is stressed by fear, hunger, illness or other aversive situations, these attachment and caregiving behaviours can *only* be observed in a stressful situation (12). Questionnaire or interview measures cannot, therefore, measure attachment - especially in early childhood, when children do not have adequate cognitive and language abilities to allow meaningful self-report (1).

In developing our new tool, we focused on early to middle childhood when interventions can be highly effective (13) and most children are in school, allowing widespread data collection for public health or epidemiological purposes. We developed a “computer game” version of the Manchester Child Attachment Story Task, called CMCAST (Minnis et al., 2010), in which the child interacted with a dolls-house as depicted on a laptop computer screen and an avatar of the child-doll and mummy-doll which he/she could move around on the screen to complete the story. The rater could then view a short animation of each story and see a cut-in video of the child as s/he told the story (see supplementary Figure).

We compared the CMCAST with the doll-play original MCAST by randomly allocating the order in which children to received CMCAST and MCAST six weeks apart, and also conducted a feasibility study on the self-administration of the CMCAST in small groups in school classrooms (14). We demonstrated that the CMCAST could be administered to school-age children with minimal adult supervision and in a classroom setting with groups of up to five children simultaneously. We were forced to pare down the CMCAST rating system to five simple elements: *child engagement* (i.e. was there a visible change in child facial expression at the crisis point of the story stem?); *child doll attachment behaviours* including proximity-seeking between child doll and mummy doll; *caregiving behaviours* from the mummy doll as depicted in the child’s story; *resolution of the attachment stress* as depicted in either the child’s story or during the post-story prompts and finally *exploratory behaviour* as depicted in the child’s story. This was a much simplified rating system compared to the more than 20 items required to classify attachment and associated features in the MCAST. Despite this, our study resulted in an almost identical distribution of attachment classifications and was as reliable as the MCAST doll-play original (14).

The efficiencies gained and the cost savings were, however, still not enough of an advance over the gold standard tools to justify production of the game. CMCAST raters still needed to attend several days of training in administering and rating the MCAST and had to work hard over several months to achieve rating reliability of both the MCAST and CMCAST. It was therefore clear that if a computerised easy-to-administer attachment measure were to be scalable and have a useful impact on attachment research and clinical practice, then automatic rating would be the next important step forward.

Simultaneously with the development of the CMCAST, we were developing an observational tool to help clinicians observe indiscriminate behaviours in the clinic waiting room. Findings showed that in an unfamiliar and therefore stressful clinic setting, typically developing children stayed very close to their parents (15) seeking for or maintaining proximity. During the SSP and in clinic waiting rooms, and of doll behaviour in doll-pay MCASTS or on the computer screen, secure attachment seemed to be characterised by *smooth* and fairly rapid movement to reduce the distance between child and mummy (doll). Movement is now recognised as a key element of human interaction and an important substrate of communicative development from before birth (16). Problems in the subtleties of body movement, such as in autism, can impede social interaction (17). The importance of movement in attachment patterns has been touched on in psychotherapeutic research (18) and practice (19), but has been barely investigated in early/middle childhood attachment research (20), probably due to the technical challenges of doing so.

Around the same time, the new computer science of “Social Signal Processing” was also being launched, asserting that “the ability to understand and manage social signals of a person we are communicating with is … a facet of human intelligence…[that is] indispensable and perhaps the most important for success in life” (21). Since then technological advances in computing science, including the development of small sensors (22) and machine learning algorithms for analysing human interactions (23), have allowed this field to move forward significantly.

We engaged some of the new advances in social signal processing in an attempt to develop a new computer “game” for measuring attachment, including movement sensors embedded in dolls and machine learning algorithms with a view to an automatic rating system (24). We have called this new computerised version of the Manchester Child Attachment Story Task (MCAST) the School Attachment Monitor (SAM), because we hope it can eventually become an efficient, low-cost way of measuring attachment in young school-age children that can be used in school classrooms for screening/mapping purposes or for data-gathering in epidemiological research.

### Our research questions were

1. Can acceptable inter-rater reliability be achieved for manual rating of SAM?
2. Is there good agreement on two-way and four-way attachment classifications between SAM and the MCAST doll-play original when manually rated by raters trained to reliability on MCAST rating?
3. For SAM, is there good agreement between a machine learning algorithm and the manual classification of secure versus insecure attachment?

## Methods

The study received ethical approcal from the University of Glasgow College of Science and Engineering Ethics Committee and the governing bodies of the local councils responsible for participating schools.

### Development of SAM

SAM was developed with close reference to the development of the Computerised MCAST, learning from design mistakes/successes with that instrument (14). Simplicity in design (a dolls-house with four rooms, no stairs and only furniture necessary for the stories and simple dolls with humanoid characteristics but no attempt at realism) aimed to a. avoid over-stimulating the child, b. facilitate the projective identification of the child with the child doll and c. encourage translation of the inner world into the play.

### Stakeholder workshop 1

A design process workshop was held with 13 typically developing children aged between 5 and 10 (7 boys; 6 girls) recruited through the personal network of the authors. The aim was to test whether children were able to engage in representational attachment story-telling on a two-dimensional surface as effectively as the traditional three-dimensional MCAST doll’s house. Children responded to up to 2 vignettes each from the MCAST using either the traditional doll’s house with dolls or on a twodimensional fuzzy felt mat as shown in Figure 1a. The order of version presentation was counterbalanced so that approximately half the children experienced doll’s house version first and vice versa. Their responses were video recorded. To our surprise, children of this age were perfectly capable of a) understanding and using a two dimensional plan version of the dolls-house in which a simple fuzzy felt mat was divided by a fuzzy felt cross into four rooms and b) of having a story stem told to them on the screen, then completing the story on the dolls-house mat, rather than on the computer screen.

**Figure 1a.**
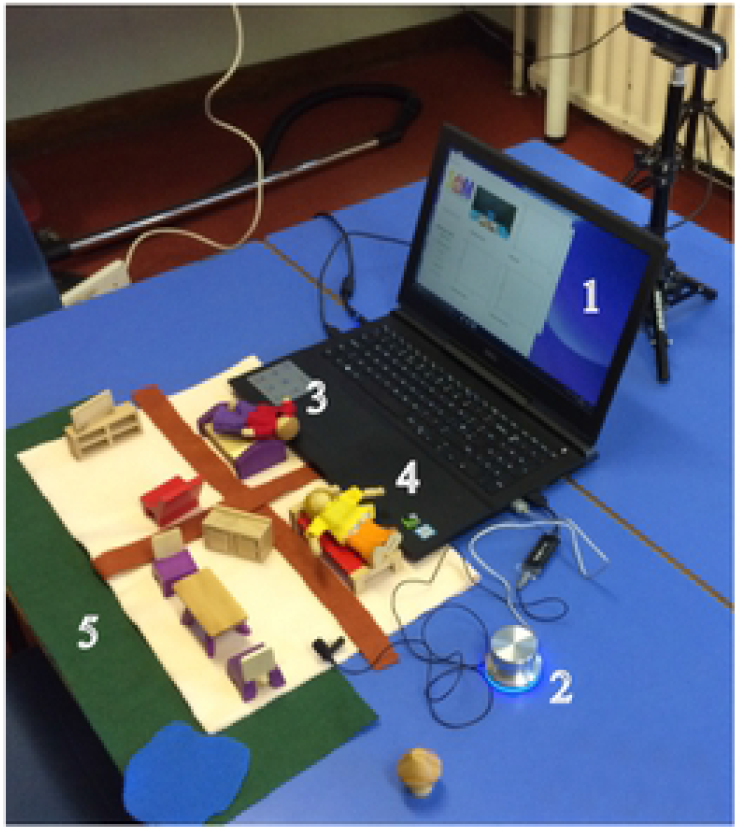
the components of SAM. (above left): shows the system as it appears to the users. The computer screen (**1**) in the picture) displays the videos with actors guiding the child through the MCAST, the button (**2**) allows the users to signal the system that they have completed an MCAST step, the dolls (**3** and **4**) and the toy house (**5**) allow the users to complete the story stems.

The use of the two-dimensional fuzzy felt mat for the dolls-house allowed us to embed tiny movement sensors into actual dolls (see Figure 1b). SAM was more like the MCAST original than the CMCAST had been in that children could manipulate both “mummy doll” and “child doll” simultaneously using real dolls (rather than screen dolls), could exploit the full range of three dimensional doll movements and therefore, like in MCAST, could fully express themselves. Like CMCAST, the entire doll and child action was videotaped with the child’s face and hands in the frame. Our administrating laptops were equipped with a specialised video camera which recorded children’s facial and upper body image as well as tracking depth to allow calculation of the distance between the dolls during their story completion. From the user perspective, these modifications simplified SAM set-up compared to CMCAST since story stems could potentially be delivered on any school computer (laptop or PC) installed with our SAM software, and the only other specific “kit” required would be the dolls-house mat, furniture, computerised dolls and the webcam-like camera with depth sensor. These are light and highly portable compared to the MCAST original set-up.

**Figure 1b.**
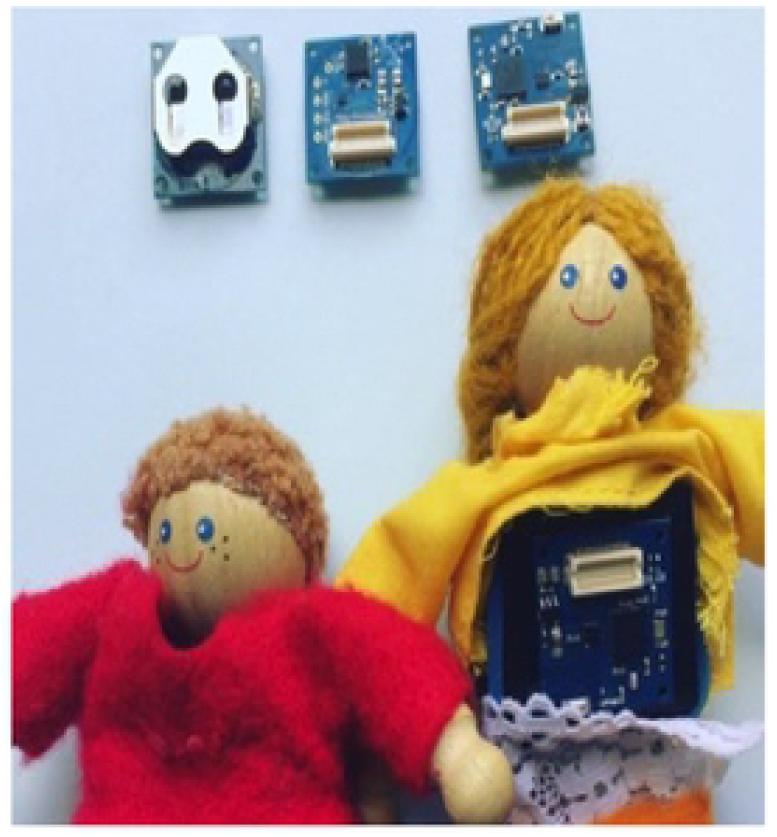
(above right): An image of the dolls that we have designed and installed with sensors inside to enable administration and machine-learning based ratings.

The five MCAST story stems (i.e., Breakfast, Nightmare, Hurt knee, Illness and shopping) were presented via a recorded performance by a professional actor who acted the beginning of the vignettes using a doll’s house. Further details are available in Vo et al. (24).

### Stakeholder workshop 2

After the *Phase 1* data collection (see below and Figure 2) a second design workshop, with the same 13 children, aimed to address any system errors. From our Phase 1 experiences, we foresaw certain errors that children might encounter during their use of SAM. These were simulated and embedded in a live puppet show with one of the researchers acting as “a puppet who needs help while playing the SAM computer game”. Children were asked to imagine a solution so that the puppet would not come across the problem again and to create a soft dough model or drawings to represent their ideas. Solutions offered included a push button with multi-functionality to communicate with SAM (see Figure 1a, element 2). In the final SAM model, children pressed this button to “talk” to the system at various points, e.g. when they have finished telling their story and are ready to move on, affording the child user a sense of agency and autonomy, essential for optimal attachment measurement in this age group. Further details are reported in (25) and the final set up is shown in Figure 1a and b (24). The overall attachment classification for SAM did not differ depending on which version of the prototype was administered: (Spearman’s rho (116) = 0.065, N.S.).

**Figure 2.**
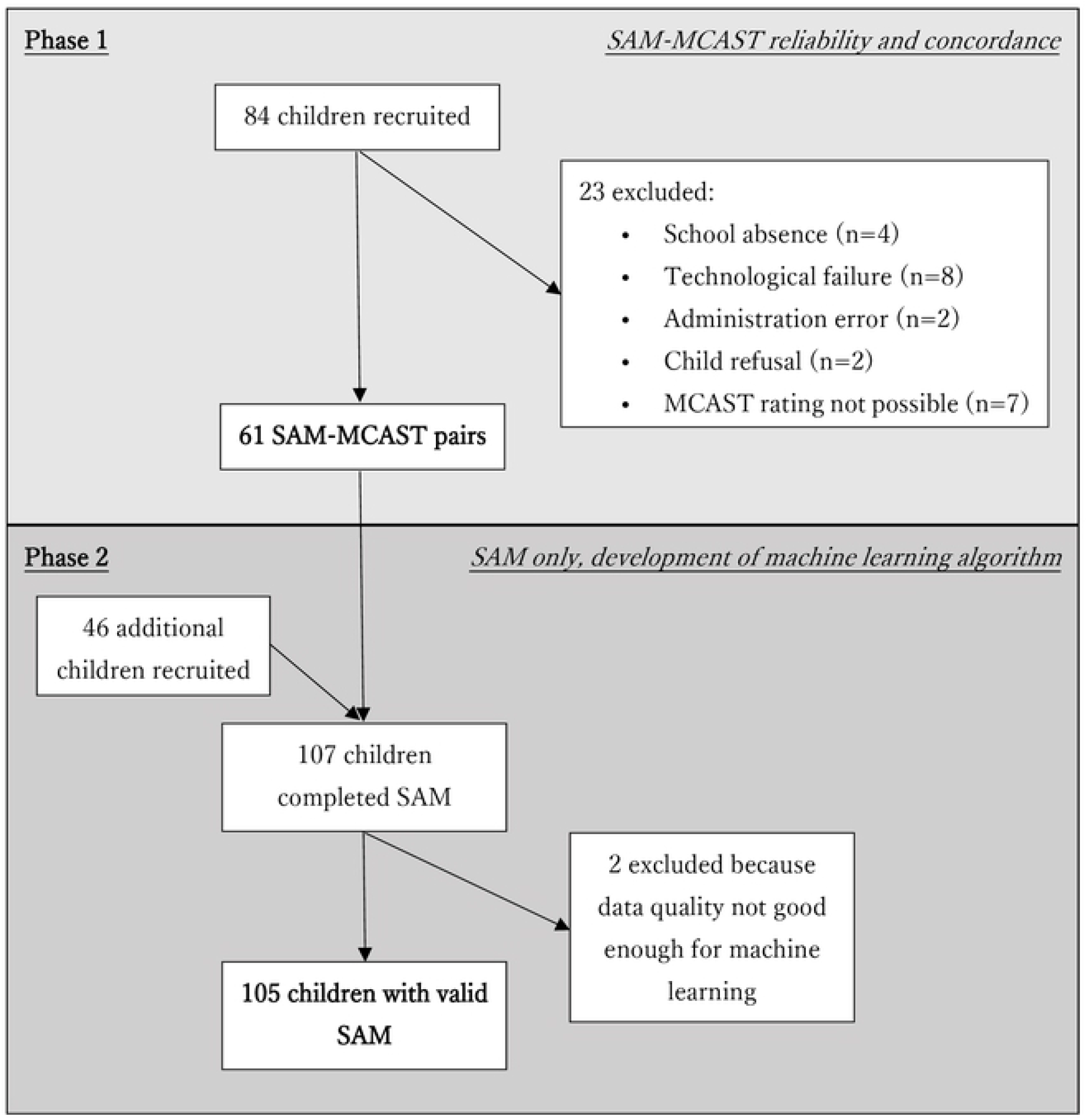
Data collection Phases 1 and 2

### Testing of SAM

Testing of SAM was conducted in two phases: *Phase 1* in which manual SAM ratings were compared with manual MCAST ratings and *Phase 2* in which an automatic SAM rating system was developed and tested against the manual SAM ratings.

### Data Collection for Phases 1 and 2

We approached schools in local councils in which the demographic background included wealthier and more materially deprived and both urban and rural areas of the West of Scotland. The Scottish Index of Multiple Deprivation (SIMD) decile, for the school addresses, ranged between 4 and 9 where the lower the value, the higher the level of deprivation for the area.

We worked with relevant staff members at each participating school to obtain consent from parents and carers of children in eligible age range, which corresponded to Years 1 to 4 (P1-4) in the mainstream primary education under the UK system. Information packs were given to the children by the school offices to take home to pass onto their families, as part of an established system for sending letters and homework to and from homes. Our information sheet explained the main aims and requirements of the study for adults (parents and carers) as well as in simple language with pictures for children, so that families could discuss participation with their children at home. Questions regarding participation and any concerns were addressed directly to the research team and families willing for their child to participate were asked to return signed consent forms by a specified date. Consent rates across the 5 schools ranged between 30 to 40%. Only children with signed parental consent were included in the study.

For Phases 1 and 2,130 children aged 5-9 (67; 51% girls) were recruited from the first 4 year groups of 5 mainstream primary schools in 3 local councils (Figure 2). For data security reasons, schools were unwilling to give exact dates of birth of the children but the distribution across school grades and age range/gender balance for each grade are summarized in Table 1.

**Table 1.** School grade, gender and age spread (%) of participating children

| *School Grade* | **Primary 1**<br>**(age 5-6)** | **Primary 2**<br>**(age 6-7)** | **Primary 3**<br>**(age 7-8)** | **Primary 4**<br>**(age 8-9)** | **Total** |
| --- | --- | --- | --- | --- | --- |
| **Number of children (%)** | 28<br>(21.5%) | 46 (35.4%) | 35(26.9%) | 21 (16.2%) | 130 |
| **% female** | 43% | 52% | 49% | 57% | 51% |

### Phase 1 – SAM – MCAST reliability and concordance

This phase addressed research Questions 1 and 2:

- Can acceptable inter-rater reliability be achieved for manual rating of SAM?
- Is there good agreement on two-way and four-way attachment classifications between SAM and the MCAST doll-play original when manually rated by trained raters?

Given our previous success with reliability and concordance comparing the CMCAST to the MCAST (14), we predicted that the new SAM system would produce data of good enough quality to allow reliable rating by trained assessors. Further, we hypothesised that the ratings across the two measures (MCAST and SAM) would show an acceptable concordance with each other as regards secure versus insecure classifications and organised versus disorganised classifications.

Each child was asked to use SAM or MCAST at least six weeks apart and data collection was organized so that similar numbers of children used MCAST first or SAM first. In Phase 1, 84 children were recruited and 61 had rateable data on both MCAST and SAM (see Figure 2).

### Manual Rating

Three raters conducted the initial manual ratings of all 61 SAM and 61 MCAST cases. One of the raters (MR) acted as a coordinator for assigning cases between the 3 raters to minimize rating bias, taking care to avoid any rating of sessions where the rater had also acted as the administrator.

As described in Figure 3, for training and quality control purposes, all 61 Phase 1 SAM cases were independently double-rated by at least two raters, then if necessary discussed in consultation with a third expert rater to agree a shared rating. Twenty percent (i.e. n=12) of the 61 MCAST cases were independently re-rated by our expert rater (SDF). After the initial training period, SDF was consulted if a rater encountered difficulties in making a judgement, if cases showed signs of attachment disorganisation or where data quality questioned whether the case was rateable.

**Figure 3.**
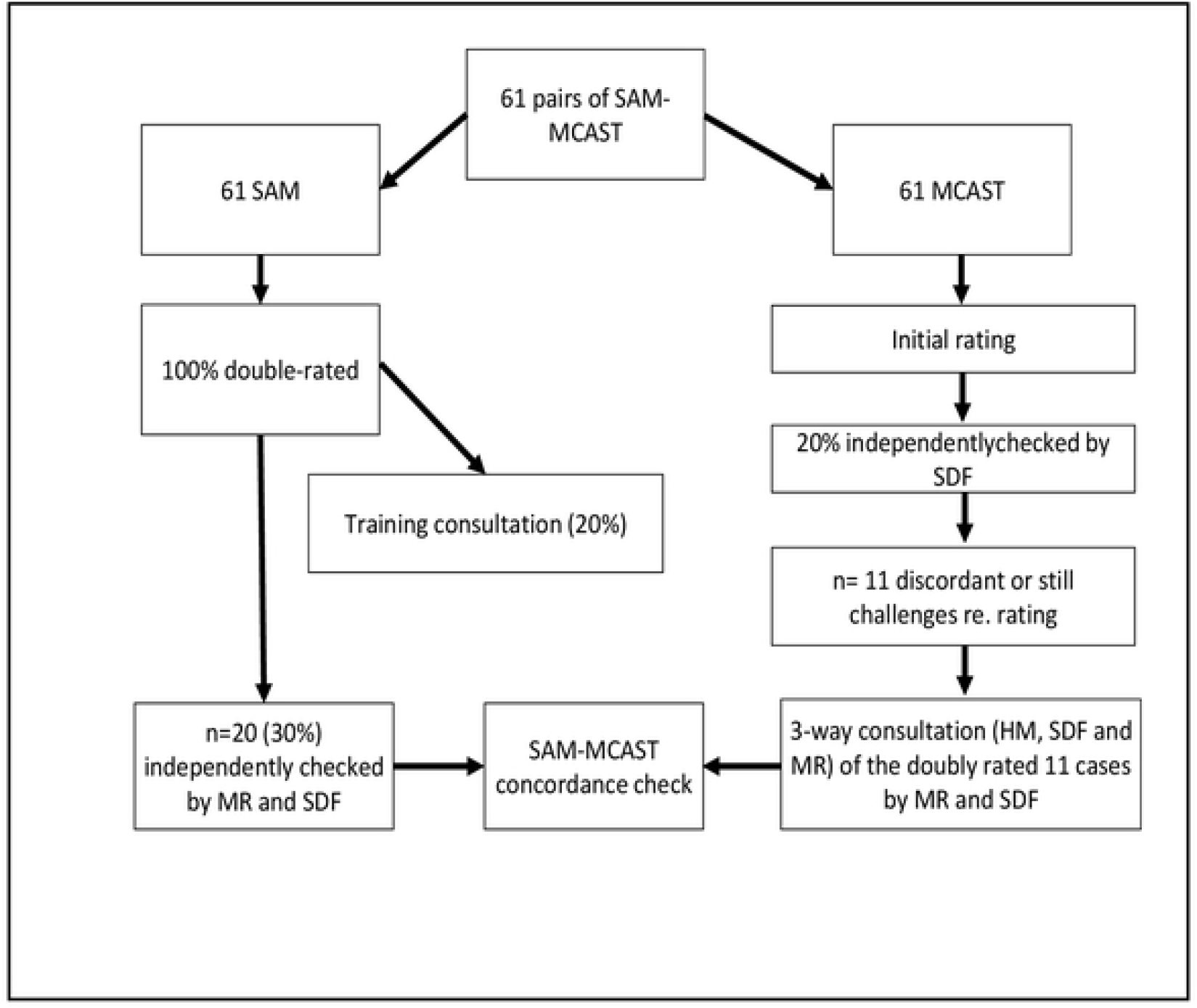
Phase 1 rating and resolution process including role of expert rater

After first rating, 11 (18%) of SAM - MCAST pairs showed significant discordance or were still challenging to rate. Since these ratings had been conducted while the team of raters was inexperienced, SDF conducted an independent re-rating of all 11 SAMs and all 11 MCASTs, never rating the SAM/MCAST for the same child on the same day. All of these ratings were then conferenced with another expert rater, HM. Throughout this second rating process, MR kept details of initial SAM and MCAST scores separately, to ensure ratings were unbiased and at no time was the SAM and the MCAST of any child discussed simultaneously during the same conference. During these conferences, each of three raters (HM, MR and SDF) examined each video and made an independent rating which was then discussed referencing the MCAST rating manual, version 26, throughout.

On average each rater was able to rate each case within 1.5 to 2 hours for MCAST, after the initial training period, and within an hour for each SAM case. We estimate that the entire manual rating process, including the ratings of all raters and the conferences, amounted to nearly 500 hours.

### Phase 2 – Development of Machine Learning Algorithm for SAM

This phase addressed research Questions 3:

- For SAM, is there good agreement between a machine learning algorithm and the manual classification of secure versus insecure attachment?

Having established the SAM-MCAST concordance in Phase 1, an additional 46 cases of SAM data were collected in order to have a sufficient sample size for the development of the machine-learning (ML) algorithm. Two of these 46 cases were unfit for ML analysis due to data quality. Together with the 61 SAM cases from Phase 1, the ML development reported here was based on a total of 105 sets of SAM data (see Figure 2). In parallel, and independently from, the manual ratings, automated ratings were made as described in detail in Roffo et al (26). In brief, the movements or “pose” of the child were analysed using OpenPose, a robust algorithm that encodes both position and orientation of human limbs in realtime from video (27). Next, OpenPose maps the coordinates of the child’s joints (See Figure 4). For SAM, the focus was on the position of the hands because of our hypothesis that the way the participants move the dolls, held in their hands, to complete the story stems is crucial for classification. We therefore focussed on the following features: *Hand position; Distance between the hands; Hand speed; Hand acceleration; Hand 1D trajectory* (i.e. placing of the hands with respect to the width of the video image); *Hand presence* (i.e. the proportion of frames in which each of the two hands is present, giving information on the child’s tendency to move one doll more than the other).

**Figure 4:**
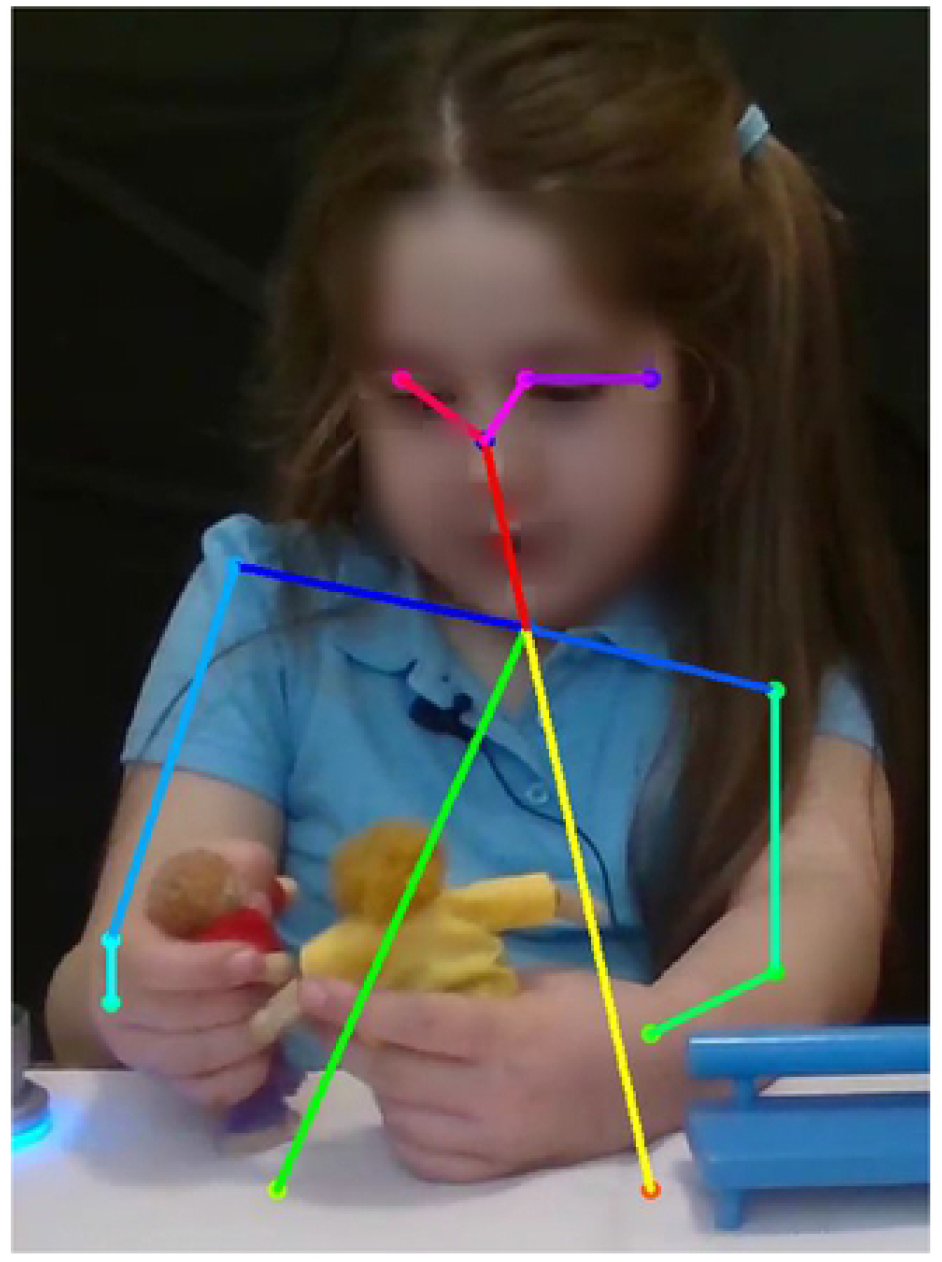
Production of a generic frame in a video using OpenPose software

The resulting preliminary algorithm was trained using the “Leave-One-Child-Out” protocol i.e. by training the algorithm with all children except one and then testing it with the only child left out. At each iteration, a different child is left out for testing and the process is therefore repeated 105 times. This allows use of the entire dataset while ensuring rigorous independence between the training and test material. The attachment classification is assigned using a “majority vote” procedure where, for each child, 500 non-overlapping video frames contribute to the decision and the child is assigned to the attachment classification most frequently extracted from these frames.

The automatic classification was then compared to the classification from the detailed manual SAM rating conducted by trained MCAST raters.

## Results

Preliminary analyses with a series of chi-square tests showed that gender, school year (age) or school attended did not differ significantly for the security or organization of attachment for either of the measurement platforms.

### Research Questions

- Is inter-rater reliability similar for both the MCAST and SAM?

#### Inter-rater realibility for SAM

the finalised ratings across the twice-rated SAM cases showed an agreement rate of 70% (Cohen’s kappa .434) at the level of four-way attachment classifications; 77.6% (Cohen’s kappa .49) for secure-insecure classifications and 95% (Cohen’s *kappa* .64) for organised-disorganised classifications (see appendix 1).

#### Inter-rater reliability for MCAST

Of the 12 cases (20%) double-rated MCASTs, there were 3 discrepancies, therefore the MCAST data showed similar four-way inter-rater reliability (~75%) compared to SAM ratings at the four-way classification level.

### Research Question 2

- Is there good agreement on two-way and four-way attachment classifications between SAM and the MCAST doll-play original when manually rated by trained raters?

As can be seen in Table 2, agreement between SAM and MCAST was excellent: secure versus insecure Cohen’s *kappa* .73 (89% agreement, discordance n= 7); organised versus disorganised Cohen’s *kappa* .78 (97% agreement, discordance n= 2) and four-way agreement Cohen’s *kappa* .73 (86% agreement, discordance n=8). For cases on which SAM:MCAST ratings agreed (in bold in Table 2 below), 39 (73%) were secure, 2 (4%) insecure avoidant, 8 (15%) insecure ambivalent, and 4 (8%) insecure disorganised.

**Table 2.** Inter-measure agreement across MCAST and SAM (N=61).

|  | **MCAST ABCD classifications** |  |  |  |
| --- | --- | --- | --- | --- |
| **SAM ABCD**<br>**Classifications** | *B-Secure* | *A- Insecure* | *C-Insecure* | *D-Insecure* |
|  | *B-Secure* | *Avoidant* | *Ambivalent* | *Disorganised* |
| *B-Secure* | **39** | 2 | 1 | 0 |
| *A Insecure*<br>*Avoidant* | 0 | **2** | 1 | 0 |
| *C Insecure*<br>*Ambivalent* | 2 | 0 | **8** | 0 |
| *D Insecure*<br>*Disorganised* | 2 | 0 | 0 | **4** |
| Total (N=61) | 43 | 4 | 10 | 4 |

*\*Cells which ratings are in agreement, are signified in bold.*

### Research Question 3

- For SAM, is there good agreement between a machine learning algorithm and the manual classification of secure versus insecure attachment?

The overall agreement between the manual rating and the automatic rating for secure versus insecure attachment was 82.8%, i.e. for 87 of the 105 children, the same attachment classification was made with the automatic rating system and the manual rating system. Children with secure patterns were identified with greater accuracy than insecure children.

## Discussion

This study has shown that, using modern sensors and machine learning technology, it has been possible to develop the School Attachment Monitor (SAM) with the following features:

- Simple administration and portability compared to the MCAST doll-play original.
- Manual rating that is similarly reliable to the MCAST doll-play original.
- An attachment distribution, using manual rating, that has good concordance with the MCAST and gives a similar distribution of four-way attachment classifications as the international literature (28).
- Most importantly, a machine learning algorithm gives an accurate classification of attachment security versus insecurity compared to manual ratings by trained MCAST raters.

The aim of our research programme, from its inception over a decade ago, was to develop a quick and easy measure of attachment that can be used in largescale public health monitoring or epidemiology. SAM has the potential to achieve that aim. Its administration is simple and the equipment required is cheap and portable. We would envisage that, for data collection purposes, schools in which there are consenting participants would be given a SAM kit for the day and the minimal research support required to ensure appropriate set up of the equipment. Throughout the day, participating children would take their turn to use SAM. From the child’s perspective, this is usually an enjoyable task that takes around 20 minutes. Data could be collected in this way from several school classrooms in a week – or from even more classrooms if simultaneous data collection using more than one SAM set-up were conducted. This could allow largescale data collection (e.g. for tens of thousands of children) to take place in a relatively short space of time and with minimal resources.

We would envisage SAM being used for largescale mapping of attachment patterns in groups of children, not for individual attachment classifications. If individual attachment measurement is required – for example as part of a clinical assessment – then we would recommend that SAM might be used as a first-stage preliminary or screening assessment, followed by a more detailed assessment using a face to face measure such as the doll-play MCAST. Even then, a recent systematic review of attachment measures has recommended that assessment of a child’s attachment pattern should never be done (especially for court reports) using a single measure, even if a gold standard measure, since no existing attachment measure has perfect psychometric properties when used alone (1). Similarly, we would argue that even gold standard attachment measures are not appropriate, as single measures, for population screening. Insecure attachment classifications are not, in and of themselves, abnormal and are regarded by many as positive adaptations to less than optimal environmental circumstances (29). On the other hand, insecure attachment is associated with increased risk of psychopathology of various types. There might therefore be potential for SAM, in future, to form part of an early, school-based screening programme to identify children at risk of psychopathology, alongside other simple screening instruments, such as the Strengths and Difficulties Questionnaire (30). If this were to be realized, such a screening programme would require careful development and testing.

Our study has certain limitations: despite achieving a good range of age gender and school deprivation category, the low recruitment rate means our study sample is unlikely to be representative of the general population, so the distribution of attachment classifications should be viewed with caution. In order to realise full potential of the SAM technology and to enable large-scale attachment studies, further technological development will be required - most critically, development of an automatic rating system which affords live rating of sessions for instant feedback. It will be important to explore the role of attachment to multiple caregivers (mother, father, and others) and the interplay between attachment patterns and other developmental (including genetic) factors. It is also critical that further testing will articulate the remit and role of SAM beyond typical development, such as in the context of adverse childhood experiences (ACEs) or developmental conditions, including neurodevelopmental disorders.

## Conclusions

We have developed a new tool for measuring attachment at the population level, which has good reliability, produces a similar distribution of attachment classifications compared to a gold-standard attachment measure and has the potential for automatic rating.

## Key Points

### What’s Known

No quick and easy measures of attachment exist that could aid clinical work or large population studies.

### What’s new

– A new attachment measure, the School Attachment Monitor, is a computer game for 5-9 year old children with simple, cheap administration. It is as reliable in classifying attachment patterns as the MCAST doll-play original and produces a distribution of attachment patterns reflecting the international database.

- A machine learning algorithm had more than 80% accuracy in classifying secure versus insecure attachment.

### What’s relevant

SAM is a quick and easy attachment measure with potential for automatic rating of attachment patterns using machine learning. This could allow largescale attachment measurement useful for public health monitoring and epidemiology.

## Acknowledgements

This research is funded by EPSRC grant EP/M025055/1. With thanks to children and schools for participating and working with the research team.

